# Predicting Moisture Content During Maize Nixtamalization Using Machine Learning with NIR Spectroscopy

**DOI:** 10.1101/2021.05.19.444884

**Authors:** Michael J. Burns, Jonathan S. Renk, David P. Eickholt, Amanda M. Gilbert, Travis J. Hattery, Mark Holmes, Nickolas Anderson, Amanda J. Waters, Sathya Kalambur, Sherry A. Flint-Garcia, Marna D. Yandeau-Nelson, George A. Annor, Candice N. Hirsch

**Author notes:** Corresponding Author: Candice N. Hirsch.

## Abstract

Lack of high throughput phenotyping systems for determining moisture content during the maize nixtamalization cooking process has led to difficulty in breeding for this trait. This study provides a high throughput, quantitative measure of kernel moisture content during nixtamalization based on NIR scanning of uncooked maize kernels. Machine learning was utilized to develop models based on the combination of NIR spectra and moisture content determined from a scaled-down benchtop cook method. A linear support vector machine (SVM) model with a Spearman’s rank correlation coefficient of 0.852 between wet lab and predicted values was developed from 100 diverse temperate genotypes grown in replicate across two environments. This model was applied to NIR data from 501 diverse temperate genotypes grown in replicate in five environments. Analysis of variance revealed environment explained the highest percent of the variation (51.5%), followed by genotype (15.6%) and genotype-by-environment interaction (11.2%). A genome-wide association study identified 26 significant loci across five environments that explained between 5.04% and 16.01% (average = 10.41%). However, genome-wide markers explained 10.54% to 45.99% (average = 31.68%) of the variation, indicating the genetic architecture of this trait is likely complex and controlled by many loci of small effect. This study provides a high-throughput method to evaluate moisture content during nixtamalization that is feasible at the scale of a breeding program and provides important information about the factors contributing to variation of this trait for breeders and food companies to make future strategies to improve this important processing trait.

**Key Message:** Moisture content during nixtamalization can be accurately predicted from NIR spectroscopy when coupled with a support vector machine (SVM) model, is strongly modulated by the environment, and has a complex genetic architecture.

## INTRODUCTION

Maize is the largest crop produced in the United States. In 2019, 13.6 billion bushels of maize were produced, which was valued at 52.3 billion dollars (United States Department of Agriculture 2019). Most of the maize grown in the United States is used for either livestock feed or ethanol production, with very little being used for direct human consumption (United States Department of Agriculture 2019). As a result of this limited relative acreage, the research and development efforts for food-grade corn varieties has also been limited. Typically, food-grade maize hybrids are high quality, high test weight maize (S.O. Serna-Saldivar et al. 1993) that is selected out of the existing No. 2 yellow dent germplasm (Holmes et al. 2019). This narrow selection process makes it difficult to make gains for specific traits of importance to food-grade maize products, such as moisture content during nixtamalization for masa-based products.

Nixtamalization is a thermal-alkaline cooking process that humans have been performing for centuries. The primary purpose of nixtamalization is to remove the kernel pericarp and soften the grain to facilitate grinding (Santiago-Ramos et al. 2018). During nixtamalization, grain are added to an aqueous alkaline solution, usually from lime (Ca(OH)_2_), and heated to a temperature above starch gelatinization, but below the boiling point of water. It remains at this temperature for between 5 and 180 minutes, depending on the needs of the end product. After cooking, the kernels are steeped in the alkaline solution for roughly 16 hours at a temperature below the starch gelatinization temperature. The pericarp is then removed either by pressurized water or mechanical forces. The remaining part of the grain, primarily endosperm, is ground into a fine flour called masa. Many of the qualities that make the end product desirable, or undesirable, are closely related to how much water is taken up by the grain while it cooks and steeps (Ramirez-Wong, B. et al. 1994). The moisture content of masa can affect properties such as texture and oil content of the end product (Holmes et al. 2019). Compositional attributes of maize vary greatly between genotypes (Flint-Garcia et al. 2009; Renk et al. 2021), and could have a profound impact on moisture content.

Maize kernels are made up of macromolecules that are unevenly distributed throughout the kernel and can impact moisture content in various ways (Holmes et al. 2019). The first macromolecules that water interacts with during the cooking process are fiber (cellulose and hemicellulose) and protein in the pericarp, which are broken down by the alkaline solution. The alkaline solution then passes through the aleurone, which is the outermost layer of endosperm cells positioned just beneath the pericarp. The aleurone is largely unaffected by the nixtamalization process (Santiago-Ramos et al. 2018), and as such should not contribute to moisture content. After passing through the aleurone, the alkaline solution proceeds to the internal endosperm, which is primarily made up of starch and protein, but also contains fiber, lipids, vitamins, and minerals. In the endosperm, the alkaline solution causes the starch granules to begin swelling. This is restricted by the protein matrix that surrounds the starch granules (Santiago-Ramos et al. 2018). Thus, when thinking about moisture content properties it is important to consider not only the starch content and structure, but also its interaction with protein content. Lipids in the endosperm can form complexes with starch molecules as well, affecting how much moisture can be taken up during nixtamalization (Santiago-Ramos et al. 2018). Lipids are especially concentrated in the germ (i.e. embryo; Weber 1979), and due to the high temperature of cooking, and long steeping time, it is possible that some lipids from the germ could diffuse to the starch granules found in the endosperm to form absorption-restricted complexes. Since the pericarp, germ, and endosperm are the primary locations of interaction with the alkaline solution, variation in the macromolecular composition of these tissues is hypothesized to be a primary driver of moisture content.

Breeding for moisture content is difficult as it is a complex trait with a number of different compositional traits likely contributing to it. Using wet chemistry to measure all of these different traits is a time-consuming process, and a faster evaluation method is preferred. Near-infrared spectroscopy (NIR), coupled with machine learning, can greatly improve the efficiency of the evaluation process. NIR is an analytical tool that measures wavebands in the near-infrared domain of the electromagnetic spectrum (750nm-2500nm) (Aenugu et al. 2011). NIR is now routinely used to identify compounds based on the combinations of specific vibrations given off when atoms bound to hydrogen absorb and subsequently release energy in the NIR spectrum. These absorbances can be quantified, and used to develop robust equations that use a combination of wet chemistry assays and spectral values to estimate macromolecular composition (Aenugu et al. 2011).

Many statistical approaches have been implemented with NIR prediction equations (Orman and Schumann 1991), but as traits become more complex, new approaches are needed. Machine learning is a branch of artificial intelligence focused on the creation of algorithms that help computers see patterns in data that humans cannot, and is a promising approach for these complex traits (Parmley et al. 2019). There are multiple forms of machine learning including supervised, unsupervised, reinforcement, and active learning. Supervised learning models are given both the predictor and response values for a large subset of the data. The computer learns patterns in the training data and is able to make predictions on new data (James et al. 2013). Machine learning gets its power from generalization, and care needs to be taken to not overfit or underfit a model (Barratt and Sharma 2018). It is vital to train on diverse, well-collected data that limit the number of confounding factors (Roh et al. 2019). There are also ways to boost generalization in machine learning models from a computational perspective, the primary method being cross-validation (specifically k-fold and hold-out). If used properly, machine learning can be a powerful tool in the realm of data science (Ornella et al. 2012).

The ability to rapidly predict moisture content during nixtamalization is necessary to make significant improvements in food-grade corn germplasm that is sourced for masa-based products. In this study, an algorithm was developed to allow accurate, high throughput prediction of moisture content during the cooking process. Moisture content values and NIR spectra from 100 genotypes grown in replicate across two environments were used to train a machine learning model. The model was then used to predict moisture content on 4,650 plots from 501 diverse genotypes grown in replicate in five environments. This information was used to partition the sources of variation and dissect the genetic basis of the trait.

## MATERIALS AND METHODS

### Plant Material and Spectral Data Collection

A set of 501 diverse maize inbred lines (Supplemental Table 1) from the Wisconsin Diversity Panel (Hansey et al. 2011) were grown in Summer 2014 and Summer 2015 in St. Paul, MN. Plants were grown with 30-inch row spacing at ^~^52,000 plants per hectare. All plants were hand pollinated and hand-harvested at physiological maturity. A subsample of 120g of grain from each plot was ground using a Perten LM 3610 Laboratory Mill with a Perten 3170 Mill Feeder. Samples were scanned using a Perten DA 7250 NIR analyzer with 141 5nm waveband absorbances from 950nm to 1650nm (Supplemental Table 2) on the large cup setting. From this spectral data, a subset of 100 genotypes were identified using k-means clustering that maximized spectral variation in the subset of selected lines within the Perten proprietary software.

The same set of 501 diverse maize inbred lines was grown at two locations in 2016 and three locations in 2017. Each location contained two replications arranged in a randomized complete block design. These inbreds flower over approximately 14 days. Within replicates, plots were blocked into early plots (flowering at approximately 71 to 80 days after planting) and late plots (flowering at approximately 81 to 87 days after planting), and randomized within the block within the replicate. In all locations plants were grown at approximately 70,000 plants per hectare with 30-inch row spacing. Experiments were grown in St. Paul, MN, and Boone County, IA in 2016 and 2017, and Columbia, MO in 2017. Open-pollinated ears from five plants per plot were hand-harvested, hand shelled, ground, and scanned using the same procedure as described above. For a small number of plots there was insufficient seed, and for these plots 80g of grain was ground and the sample was scanned on the small cup setting. For all samples that were scanned, the waveband absorbances were exported (Supplemental Table 3) and estimated trait values for macromolecules were generated using existing Perten equations that were calibrated using Honig’s Regression for percent fat, percent fiber, percent protein, percent starch, and percent ash (Supplemental Table 3).

### Laboratory-Scale Cooking Procedure

A benchtop cook test was used to assess moisture content during the cooking and steeping steps of the nixtamalization process. Seed from the 2016 plots grown in St. Paul, MN and Boone County, IA for the 100 genotypes that were identified to maximize spectral variation were subjected to this bench top cook test. Both of the replicates grown in St. Paul, MN, and Boone County, IA in 2016 were cooked in duplicate according to the method below, which equates to ^~^800 total cooks (Supplemental Table 4). The benchtop cook test was performed on a 100g sample of grain. Grain was dried in a 27°C oven for 24 hours prior to cooking to ensure consistent moisture content. For the cooking, 2.4g of calcium hydroxide was added to 400ml of deionized water in a 1L beaker. A stir bar was placed in the bottom of the beaker under a cook basket. Raw, dried corn kernels were poured into the cooking basket. The beaker was covered with aluminum foil and placed on a pre-heated hotplate (60°C) that was then heated to 93.3°C and temperature maintained for one minute. The samples were immediately quenched with 200ml of room temperature deionized water, the cooking basket was removed from the beaker, and kernels were poured back into the beaker of nejayote and placed into a 49°C water bath to be steeped for 16 hours. After steeping, samples were removed from the bath and the pH of the solution was determined. The kernels were then poured into a large metal bowl with the nejayote. Kernels were picked up and rubbed by hand to remove pericarp and debris. After this, the samples were poured through a strainer into a 2L beaker. Kernels were then placed back into the bowl with 400ml of deionized water. Kernels were hand rubbed to remove pericarp three more times (for a total of four repetitions), each time straining and using fresh deionized water. Washed kernels were put into an aluminum tray which was zeroed on a single decimal scale in order to obtain the wet kernel weight. Two subsamples of ^~^15g of kernels from each cook (designated as Y and Z in Supplemental Tables 4 and 5) were measured into their own trays and the mass with and without the tray was documented as “Wet Kernel Weight with Tray” and “Wet Kernel Weight,” respectively. These subsamples were placed with their trays in a 103°C oven to dry for one week and were weighed again to obtain the “Dry Kernel Weight” (Supplemental Figure 1).

To determine the moisture content, equation 1 was applied for each of the subsamples that were dried. The average of the two 15 g-subsamples was taken to determine the moisture content of a sample cook, and the average of the two duplicated cooks from a plot was then taken to determine the moisture content of a given plot. Plots with too little seed to cook, and plots with missing moisture content values for all four subsamples were removed, leaving the dataset with information from 316 of the original 400 plots (2 locations x 2 replicates x 100 plots) (Supplemental Table 4).

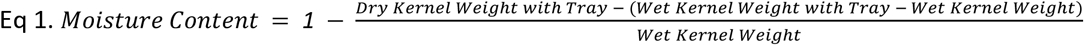

### Statistical Methods Applied to Plots of the Spectrally Diverse Genotypes

Statistical outliers were removed by one of two methods as determined by the data type. Macromolecular trait outliers were removed if the value was more than three standard deviations away from the mean of a given trait. Spectral outliers were removed based on a Mahalanobis distance (P.C. Mahalanobis 1936) of three for every other waveband to limit collinearity issues. After removing samples with missing data, only one sample was removed as an outlier. Spectral normalization was then performed by max norm, absolute value norm, euclidean norm, and standardization.

An analysis of variance (ANOVA) was conducted to dissect sources of phenotypic variation for the macromolecular traits and percent moisture content on the 316 plots of spectrally diverse genotypes using the lme4 and lmerTest packages in R version 4.0.3 (R Core Team 2020). In this analysis all factors were treated as random effects. Equation 2 shows the formula for this model.

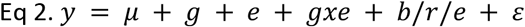

For moisture content, a second ANOVA was performed with these same 316 plots using a type II sum of squares approach with the cars package in R version 4.0.3 (R Core Team 2020) that included the macromolecular traits, spectral waveband absorbances, and cooking parameters collected during the cooking process (i.e. cook time, steep time, and pH). In this model, macromolecular traits, spectral absorbances, and cooking parameters were treated as fixed effects.

Pearson correlation coefficients between moisture content and each of the variables (NIR estimated macromolecular traits, spectral wavebands, and cooking related traits) collected on the 316 plots of spectrally diverse genotypes were determined using the cor function in R version 4.0.3.

### Machine Learning Model

Machine learning models for predicting moisture content were first generated using the NIR-estimated macromolecular traits as features, and subsequently using spectral data as features. For this analysis, the caret (Kuhn 2008) package version 6.0-86 was used in R version 4.0.3 (R Core Team 2020). The models trained on composition trait features included rpart, knn, lm, rf, svmLinear, svmPoly, and svmRadial, while models trained on spectral absorbances include pls and pcr in place of rpart and knn. For both compositional traits and spectra, the data were split into training and validation sets using two different partitioning methods, random partitioning and partitioning based on genotype, both using an 80/20 split. Ten-fold cross validation by random fold assignment was performed for models using randomly partitioned data and genotype-group assignment for models using by-genotype partitioned data. In genotype-based cross-validation, all samples of a given genotype were within either the training or testing fold for any given cross validation. This increases the generalizability of the model as it does not allow information from training on a genotype to inform the predictions on the same genotype.

Models were given a list of hyperparameter values through the training argument tuneGrid. Macromolecular-trait-based models were given a total of 5 hyperparameter values, whereas spectra based models were given a total of 141 (one for each feature). For models such as rf, pls, and pcr, where the number of hyperparameters cannot exceed the number of features, the hyperparameter value was equal to the iteration value. For models that had the ability to have near infinite hyperparameter values (such as SVM’s), the primary hyperparameter value (C in the case of SVM’s) was set based on a mathematical formula depending on the model. The linear regression model’s intercept was equal to the iteration value (i) divided by the maximum iteration value (n=141), and the support vector machine’s cost value was equal to i^2^ x 0.007. This allowed hyperparameters to be set below 1 as well as near the maximum iteration value for models such as support vector machines. Secondary hyperparameter values (only necessary for SVM’s) were not controlled, and instead given a short list of values to evaluate that included 1/2/3 and 0.001/0.01/0.1 for svmPoly’s degree and scale hyperparameters respectively, and 0.001/0.01/0.1 for svmRadial’s sigma hyperparameter.

To determine the best combination of hyperparameters, spectral normalization, cross validation methods, and partitioning variables, models were bootstrapped 100 times through all combinations of variables. Models trained on compositional traits varied by model (n=7), partitioning and cross validation method (n=2), and hyperparameter values (n=5), for a total of 70 combinations that were each bootstrapped 100 times. Spectra-based models were the same with the exception of an increased number of hyperparameters (n=141) and an addition of spectral normalization method (n=5; 4 normalizations plus the base spectra) for a total of 9,870 combinations that were each bootstrapped 100 times. Within each iteration, the seed value was set to the bootstrap iteration value (1:100), to ensure all random events occurred the same across all combinations within an iteration, but differently across iterations.

Machine learning models were evaluated on two metrics: root mean squared error (RMSE) and Spearman’s rank correlation coefficient (r_s_), with priority given to r_s_. Learning curves of the top model for both macromolecular and spectra-based predictions were created using the learning_curve_dat function in the caret R package (Max Kuhn 2020). This function was bootstrapped 100 times to reduce the effect of random sampling on the final result. The arguments test_prop and proportion were set to 0.1 and 5:20/20, respectively. All other arguments were pulled directly from the train function of the respective model. Resampling data were discarded, and an average training and testing performance was calculated by grouping the data by both training size and data type (training/testing).

### Model Validation

The best model and hyperparameter combination (linear SVM kernel, C = 71.407, trained on absolute value normalized spectra, cross validated by genotype) was then retrained using the full set of 316 plots. This retrained model was used to predict moisture content for the full set of plots grown in the 2016 and 2017 trials described above that had sufficient grain for NIR spectroscopy (n=4,650 plots; Supplemental Table 3). From these predictions, 47 plots were selected that span the range of predicted moisture content values, including nine plots that fell outside the training bounds. These plots did not include any of the spectrally diverse genotypes used to train the model. Grain from the 47 selected plots were cooked in replicate according to the methods described above (Supplemental Table 5). Moisture content of these validation samples were calculated (equation 1) and compared to the predicted value from the retrained model. R_s_ and RMSE values were calculated to summarise the model validation. For downstream analysis, any plots from the n=4,650 predicted plots that had values outside of the boundaries (n=663) of the training set (37.1-53%) were removed due to low extrapolation power of the model.

### Calculation of Feature Importance

Feature importance testing was performed following the previously proposed model reliance method (Fisher et al. 2019). For this analysis, a model was first generated from the 316 cooked plots. The model was then used to predict the 38 of 47 validation plots that fell within the training bounds (closed circles in Figure 3B) and a Pearson correlation coefficient was calculated between the predicted values and the wet chemistry values for these plots. The model was then used to predict moisture content on the 38 validation plots that were within the training bounds with a single waveband permuted at a time. Each waveband was permuted 100 times. Pearson correlation coefficients for these predictions were compared to the baseline correlation to determine the reliance of the model on each waveband.

### Statistical Analysis for Predicted Moisture Content in the Trial Plots

To determine the sources of variation in moisture content predictions, an ANOVA was run on the predictions of the full set of plots grown in 2016 and 2017. In this analysis, moisture content prediction was the dependent variable, and genotype, environment, genotype-by-environment interaction, and block nested within replication nested within environment, were the independent variables and were treated as random effects (equation 2). This was computed using the lme4 and lmerTest packages in R version 4.0.3 (R Core Team 2020). The dataset was then partitioned to only contain plots that correspond to dent genotypes defined as stiff_stalk, non_stiff_stalk, and iodent in Supplemental Table 1 (n=3,155). An ANOVA was run on this subset of plots using the same model applied to the full data set.

Heritability for predicted moisture content was determined on an entry-mean basis. Calculations of heritability were performed using the output of the mixed linear model described above for all germplasm types (n=3,987), using equation 3.

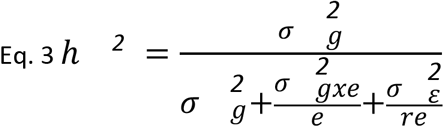

Correlations between macromolecular traits and moisture content were determined using the cor.test function in base R version 4.0.3. This was calculated for both the training (actual moisture content from the benchtop cooking protocol; n=316) and prediction (predicted moisture content; n=3,987) sets.

### Genetic Dissection of Predicted Moisture Content

Genome wide association studies (GWAS) were performed as previously described (Renk et al. 2021). Briefly, GAPIT version 3 (Lipka et al. 2012) was used to transform genomic data into numeric format, generate a genotypic map dataset, and a PCA covariates dataset (n=5). SNP data were obtained from a previous study (Qiu et al. 2021), and filtered as previously described (Renk et al. 2021). Because environment was a large source of variation based on the ANOVA described above, GWAS was performed separately for each of the five environments. The trait data were split by environment, from which BLUPs were extracted with the random effects model described in equation 4, where y = predicted moisture content. To extract the BLUPs from this model, the function ranef from the lme4 package (Bates et al. 2015) in R 4.0.0 was used. Genotypes in the BLUP dataset for each environment were filtered to only include those present in the SNP dataset.

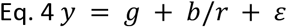

After the BLUP datasets had been created for each environment (n_MN2016_=426, n_IA2016_=442, n_MN2017_= 281, n_IA2017_= 409, n_MO2017_=412), the numeric genomic dataset was filtered to only include rows where the genotype was present in the BLUP dataset. The genomic map, filtered numeric genomic, and filtered BLUP datasets were used in conjunction to permute 30 p-values using the function FarmCPU.P.Threshold in FarmCPU version 1.02 (Liu et al. 2016) in R version 4.0.0. To run FarmCPU, the filtered BLUP, filtered numeric genomic, genomic map, covariates, and permuted p-value datasets were utilized. A reliable p-value for the FarmCPU function was determined by using the 0.05 quantile of p-values from the permuted dataset. The threshold.output was set to 1, MAF.calculate to true, method.bin to “optimum”, maf.threshold to 0.05, and maxLoop to 50.

After significant SNPs were determined in each environment, MaizeGDB (Maize B73 RefGen_v4) (Monaco et al. 2013), NCBI BLAST (Altschul et al. 1990) and Pfam (Mistry et al. 2021) were used to determine which genes the SNPs were located in or near and a putative function for each gene. MaizeGDB was used to locate and identify genes near the SNP with the Zm-B73-REFERENCE-GRAMENE-4.0 Zm00001d Gene Set and NCBI B73_v4 annotation release 102 tracks enabled. SNP distance was calculated in kilobases away from the beginning (positive numbers) or end (negative numbers) of a gene, or as zero if present within a gene. The sequence of the closest gene was obtained from MaizeGDB and inserted into NCBI BLAST (blastn=genomic sequence, blastx=cDNA sequence) and Pfam (cDNA sequence) search fields to identify putative functional annotations.

Percent variance explained by the significant SNPs as well the entire SNP set was determined through GCTA (v1.93.2 beta; Yang et al. 2011), as previously described (Renk et al. 2021). Briefly, the original SNP data (Qiu et al. 2021) was filtered for genotypes present in a given environment and formatted into a binary hapmap format using TASSEL 5 (Bradbury et al. 2007), and then saved as .ped and .map files within TASSEL 5. PLINK (v1.07; Purcell et al. 2007; Shaun Purcell) was used to create a .bed file of the binary dataset, which was then used in GCTA (v1.93.2 beta; Yang et al. 2011) to create a genetic relatedness matrix which was used in GCTA along with the moisture content predictions used in the GWAS to determine the percent variance explained by the significant SNPs and the entire SNP set in each environment.

### Code Availability

All code is publicly available on GitHub at https://github.com/HirschLabUMN/ML_Moisture_Prediction.

## RESULTS

### Variation in the Training Data

The genotypes for the training dataset were selected from a panel of 501 diverse inbred lines that encompass variation that exists in northern maize temperate germplasm (Hansey et al. 2011). This panel was grown in two locations (St. Paul, MN and Ames, IA) over two years, and grain samples were evaluated with NIR spectroscopy. In total, 100 genotypes were selected that provided maximum spectral variation across 141 wavebands (Figure 1A). This set of 100 genotypes included 67 dent genotypes, 2 popcorn/sweetcorn/flint genotypes, and 31 other genotypes. Spectral reflectance/absorbance is a product of the compositional attributes of a sample, specifically it is measuring overtone and combination bands of vibrations from hydrogen bearing groups. Thus, in selecting spectrally diverse genotypes, the genotypes should also be compositionally diverse. Indeed, the 100 spectrally diverse lines varied in macromolecular compositional traits predicted from NIR spectroscopy including percent protein, percent starch, percent fiber, percent fat, and percent ash (Figure 1B). The ranges for each trait in the training set were 5.75% to 17.14% for protein, 66.93% to 74.15% for starch, 1.18% to 2.53% for fiber, 2.85% to 6.19% for fat, 1.11% to 1.81% for ash, and 37.1% to 53.0% for moisture content. An analysis of variance (ANOVA) was performed on each of the macromolecular traits for these 100 lines grown in four environments (location-by-year combinations) (Figure 1C). Variation due to genotype, environment, and genotype-by-environment interaction vary by trait, though in all traits except starch, genotype explains the largest portion of variation.

**Figure 1.**
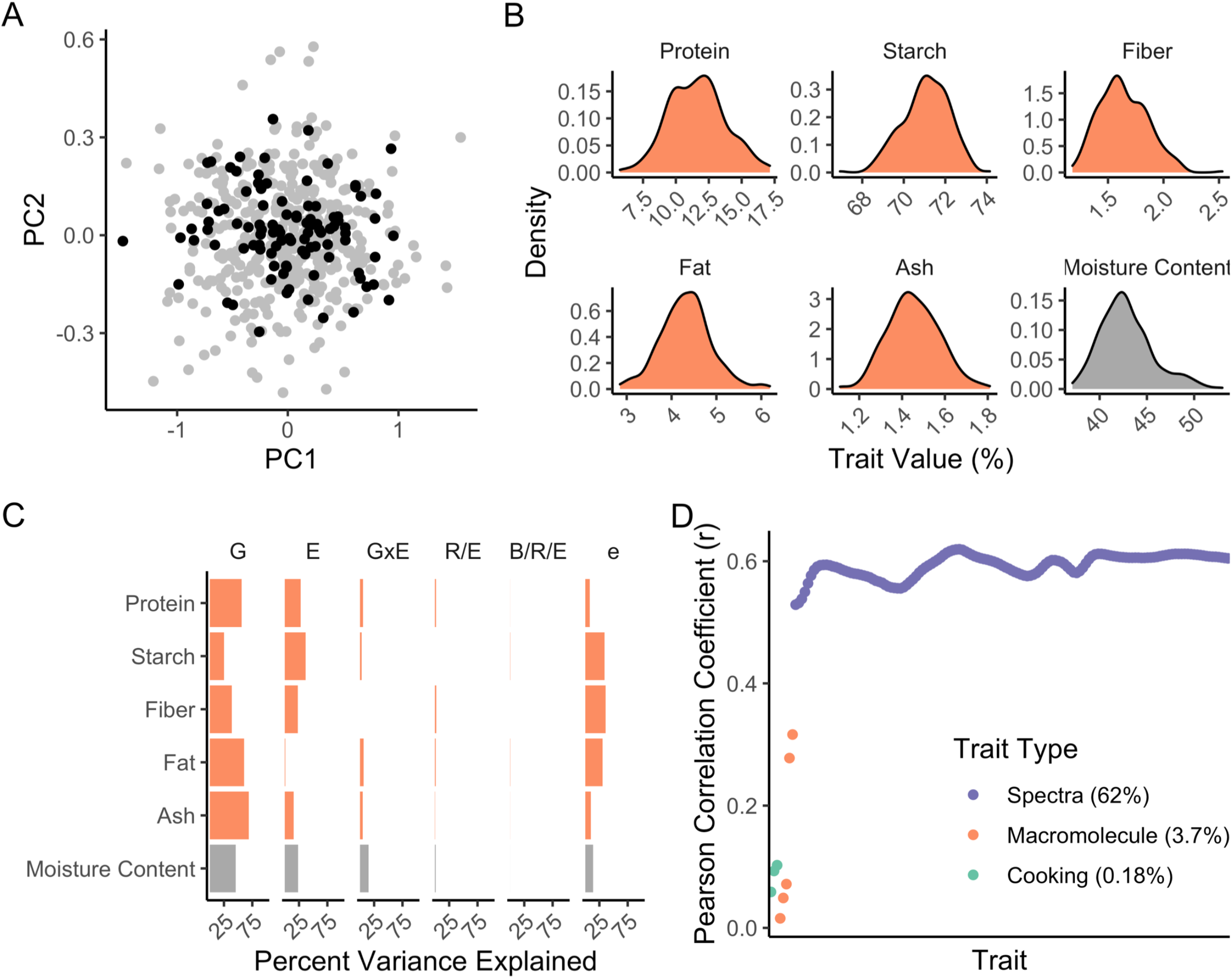
Variation observed in the training set used to develop the moisture content prediction model. A) Principal Component Analysis (PCA) plot with the full set of genotypes (gray) and the 100 selected for spectral variation (black). B) Distributions of the macromolecular traits obtained from NIR spectroscopy and moisture content from the benchtop cooking protocol for samples in the training set. C) Partitioning of variation sources among the macromolecular traits and moisture content in the training set. D) Correlation within the training set of moisture content with each of the cooking parameters, NIR-estimated macromolecular traits, and spectral wavebands. G, genotype; E, environment; GxE, genotype-by-environment interaction; R/E, replicate nested within environment; B/R/E, block nested within replicate nesting within environment; e, error.

Moisture content during nixtamalization for the plots of spectrally diverse genotypes in the multienvironment trial was measured using a benchtop cooking protocol (Supplemental Figure 1). A wide range in variation was observed for moisture content, with a minimum of 37.1% and a maximum of 53.0% (Figure 1B). Analysis of variance revealed a significant portion of the variation was explained by genotype (46%), environment (24%), and genotype-by-environment interaction (15%) (Figure 1C). Pearson correlation coefficients between moisture content and cooking parameters (cooking time, steeping time, pH), macromolecular traits (percent protein, percent starch, percent fiber, percent fat, and percent ash), and spectral bands (n=141) revealed significant correlations with fat, starch, and all 141 wavebands (Figure 1D; p-value < 0.05), though the correlations with the wavebands were substantially stronger than for the cooking parameters and macromolecular traits.

### Prediction Models Trained on Macromolecular Features

Compositional traits have more direct biological significance than spectral data and, for this reason, were first used to train prediction models for moisture content during nixtamalization. A total of 70 combinations of models (n=7), partitioning and cross validation (n=2), and hyperparameters (n=5), each with 100 iterations (total n=7,000 iterations) were trained to determine the optimal combination to predict moisture content during nixtamalization from macromolecular composition of raw kernels. Spearman’s rank correlation coefficient (r_s_) between predicted and observed values were used to assess which combination consistently generated the best model across iterations (Supplemental Table 6). Random partitioning and cross validation outperformed by-genotype partitioning and cross validation in training (Supplemental Table 6), but cross validating by-genotype performed better in testing. As such, focus was placed on models trained on by-genotype partitioning and cross-validation. The top performing combination of each model is shown in Table 1. The linear regression model (lm) with an intercept of 0.2 performed the best with an average r_s_ value of 0.487, followed by the linear SVM kernel (svmLinear; 0.486), the radial SVM kernel (svmRadial; 0.478), the polynomial SVM kernel (svmPoly; 0.474), the random forest model (rf; 0.399), the decision tree model (rpart; 0.306), and the k-nearest neighbor model (knn; 0.286). The range in performance for by-genotype models was as high as 0.066 to 0.738 for svmRadial, and on average there was a range of 0.520 (Figure 2A). Similar ranges in values across the 100 iterations were also seen for the by-random models (Supplemental Figure 2). These performances, while better than guesses, were relatively low even for the best model trained from the macromolecular trait data. A learning curve for the best model (lm) demonstrated a good overall fit based on the relative proximity of the lines near x=250 (Figure 2B), indicating the addition of more samples would not lead to major performance improvements.

**Table 1:** Performance of top model combinations.

| Model<sup>a</sup> | Normalization | Data | Hyperparameter | Hyperparameter Value | Performance<sup>b</sup> |
| --- | --- | --- | --- | --- | --- |
| lm | NA | Macro | intercept | 0.2 | 0.4870 |
| svmLinear | NA | Macro | C | 5 | 0.4866 |
| svmRadial | NA | Macro | C; sigma | 5; 0.001 | 0.4786 |
| svmPoly | NA | Macro | C; degree; scale | 5; 2; 0.01 | 0.4740 |
| rf | NA | Macro | mtry | 5 | 0.3990 |
| rpart | NA | Macro | cp | 0.02 | 0.3062 |
| knn | NA | Macro | k | 5 | 0.2860 |
| svmLinear | Absolute Value | Spectra | C | 71.407 | 0.7266 |
| pls | Base Spectra | Spectra | ncomp | 16 | 0.7264 |
| pcr | Absolute Value | Spectra | ncomp | 28 | 0.7229 |
| svmPoly | Absolute Value | Spectra | C; degree; scale | 139.167; 1; 0.1 | 0.7120 |
| svmRadial | Sum of Squares | Spectra | C; sigma | 10.647; 0.01 | 0.6038 |
| rf | Base Spectra | Spectra | mtry | 129 | 0.5830 |
| lm | Absolute Value | Spectra | intercept | 0.29078014 | 0.5566 |

<sup>a</sup>Models were partitioned and cross validated by-genotype
<sup>b</sup>Spearman's rank correlation between predicted values and values obtained from the benchtop cook test

**Figure 2.**
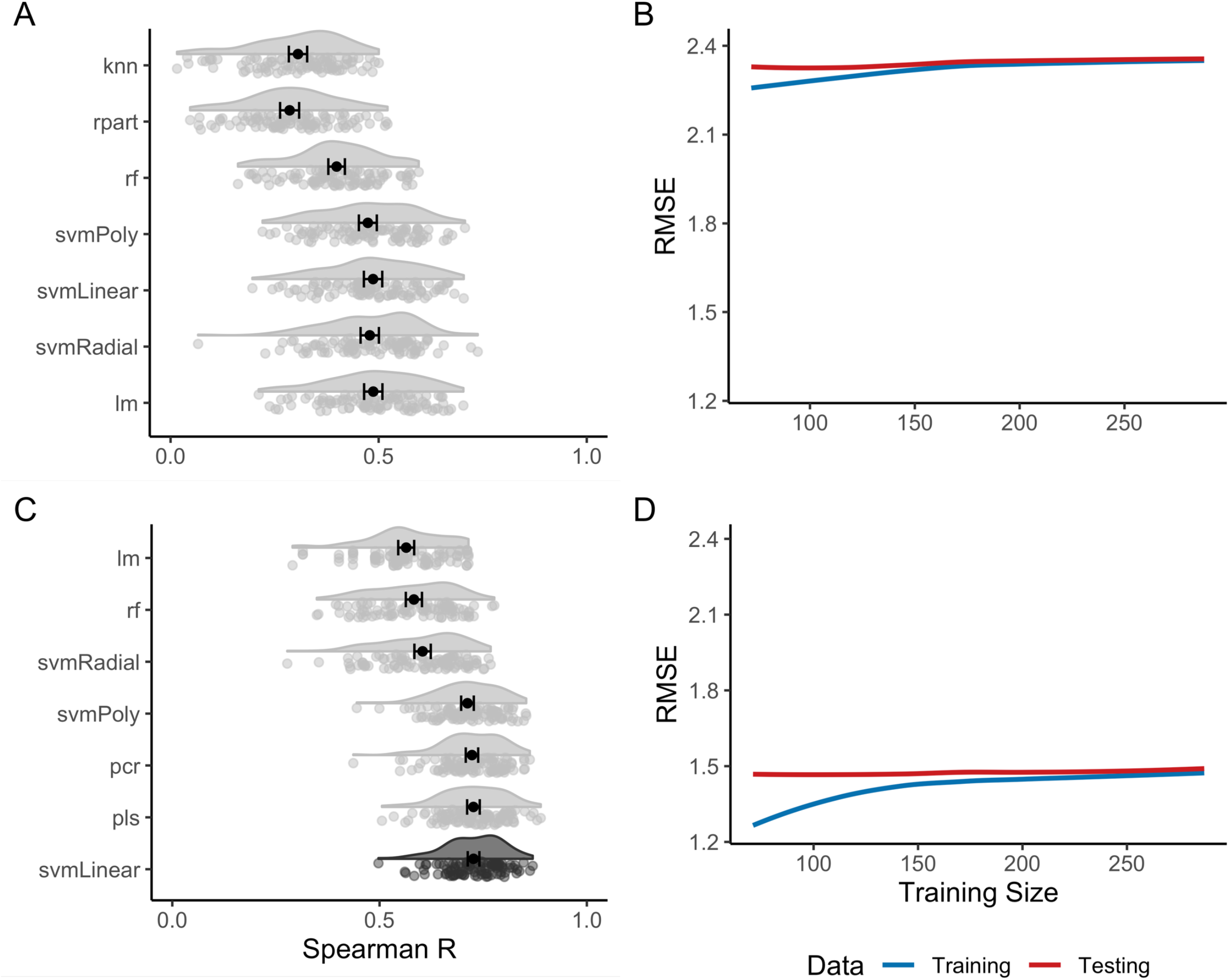
Performances of the top macromolecular and spectra-trained models. A) Bootstrapped performance of genotype-partitioned and cross-validated models trained on macromolecular traits to predict moisture content. Each data point for A and C are a single iteration of the bootstrapped model in which a variation of each model was trained on 80% of the dataset, and tested on the remaining 20%. Supplemental Figure 3 demonstrates what each point represents. B) Learning curve of the macromolecular-based linear regression model, which shows very minimal improvement from added training samples above n=150. C) Bootstrapped performances of genotype-partitioned and cross-validated model trained on spectra to predict moisture content. The dark gray distribution is the model that performed the best and was used for downstream analyses. D) Learning curve of the spectra-based svmLinear model, which shows limited improvement with training sizes greater than n=200.

### Prediction Models Trained on Near-Infrared Spectroscopy Spectral Features

The macromolecular traits used to train the models described above were predicted from NIR spectroscopy. While these macromolecular traits hold more biological meaning than the 141 wavebands from which they were generated, it is possible that there is information contained within the spectral data that is not represented within these derived compositional traits. The spectral data were more highly correlated with moisture content during nixtamalization (Figure 1D), supporting this hypothesis. Furthermore, a type II sum of squares ANOVA on moisture content as a function of cooking parameters (cook time, steep time, pH), composition (protein, starch, fat, fiber, ash), and spectral absorbances (950nm-1650nm) showed that spectra explained 62% of the variation in moisture content, whereas macromolecular composition and cooking parameters explained 3.7% and 0.18%, respectively. As such, models were developed using the spectral data directly as features to try to improve the prediction accuracy.

As with the macromolecular traits, combinations of models (n=7), partitioning and cross validation (n=2), hyperparameters (n=141), and spectral normalizations (n=5), each with 100 iterations (total n=987,000 iterations) were trained to determine the optimal combination to predict moisture content during nixtamalization from raw kernel spectral profiles (Table 1; Supplemental Table 7). Similar to the macromolecular combinations, it was found that the best training model was dependent on the choice of partitioning and cross validation. The best model during random partitioning and cross validation was the partial least squares model (pls), while the linear SVM kernel performed the best when partitioned and cross validated by genotype. The linear SVM kernel trained on by genotype partitioning and cross validation outperformed the pls model trained on random partitioning and cross validation in testing, which again suggested increased generalization from a decrease in information leakage (Figure 2C, Supplemental Figure 2). Due to this increase in generalization, we focused on models trained from by-genotype partitioning and cross validation. The top model combinations had very similar distributions of bootstrapped performances, with the top models having nearly identical average performances (Figure 2C). In general, the linear SVM kernel showed a strong predictive ability for the absolute value normalized spectral dataset as hyperparameter variations of this combination occurred 58 times in the top 60 by-genotype models (Supplemental Table 7). Ultimately, the linear SVM kernel with by-genotype partitioning and cross validation, absolute value normalization, and a hyperparameter of C = 71.407 was chosen as the final combination for subsequent predictions as it had the highest performance (average r_s_=0.726). The learning curve for this model was created in the same manner as the macromolecular model based learning curve. As with the top macromolecular model learning curve, the spectra-trained model learning curve shows little room for improvement when more samples are cooked (Figure 2D). In contrast to the macromolecular model learning curve, however, the spectra model learning curve had a ^~^40% decrease in root mean squared error, meaning the predictions were closer to the actual values obtained during cooking.

### Near-Infrared Spectroscopy Trained Model Validates Across the Training Bounds

The linear SVM kernel trained on the full data set (n=316) was used to predict moisture content for 4,650 plots that were grown across two years and three locations (5 total environments) (Supplemental Table 3). Moisture content prediction was normally distributed with a mean of 41% and a range of 27.3%to 57.3% (Figure 3A). The training bounds for the model were from 37.1% to 53.0% (filled bars in Figure 3A), and 14.3% of predictions were outside of these bounds (empty bars in Figure 3B). From this distribution, 47 plots were selected that spanned the range of predicted moisture content across the plots (tick marks under the graph in Figure 3A). These plots include 38 samples that were within the boundaries of the training data and 9 plots that were outside of these boundaries. These 47 samples were cooked using the benchtop cooking test (Supplemental Figure 1) to obtain moisture content during nixtamalization (Supplemental Table 5). Plots inside of the training set bounds were used to validate the model on an independent set of plots not used in the training set. Plots outside of the training set bounds were used to determine the model’s ability to extrapolate beyond the training set bounds. For the 38 plots that were within the training boundaries (closed circles in Figure 3B), a very high significant correlation was observed (r_s_=0.852; p-value <0.05). In contrast, the nine plots that had predictions outside of the original training bounds (open circles in Figure 3B) showed poor validation accuracy. However, the training set used in this study included samples well outside what would traditionally be accepted for moisture content in the industry (46-51% depending on the product), and the plots with extrapolated values would not be considered for advancement.

**Figure 3.**
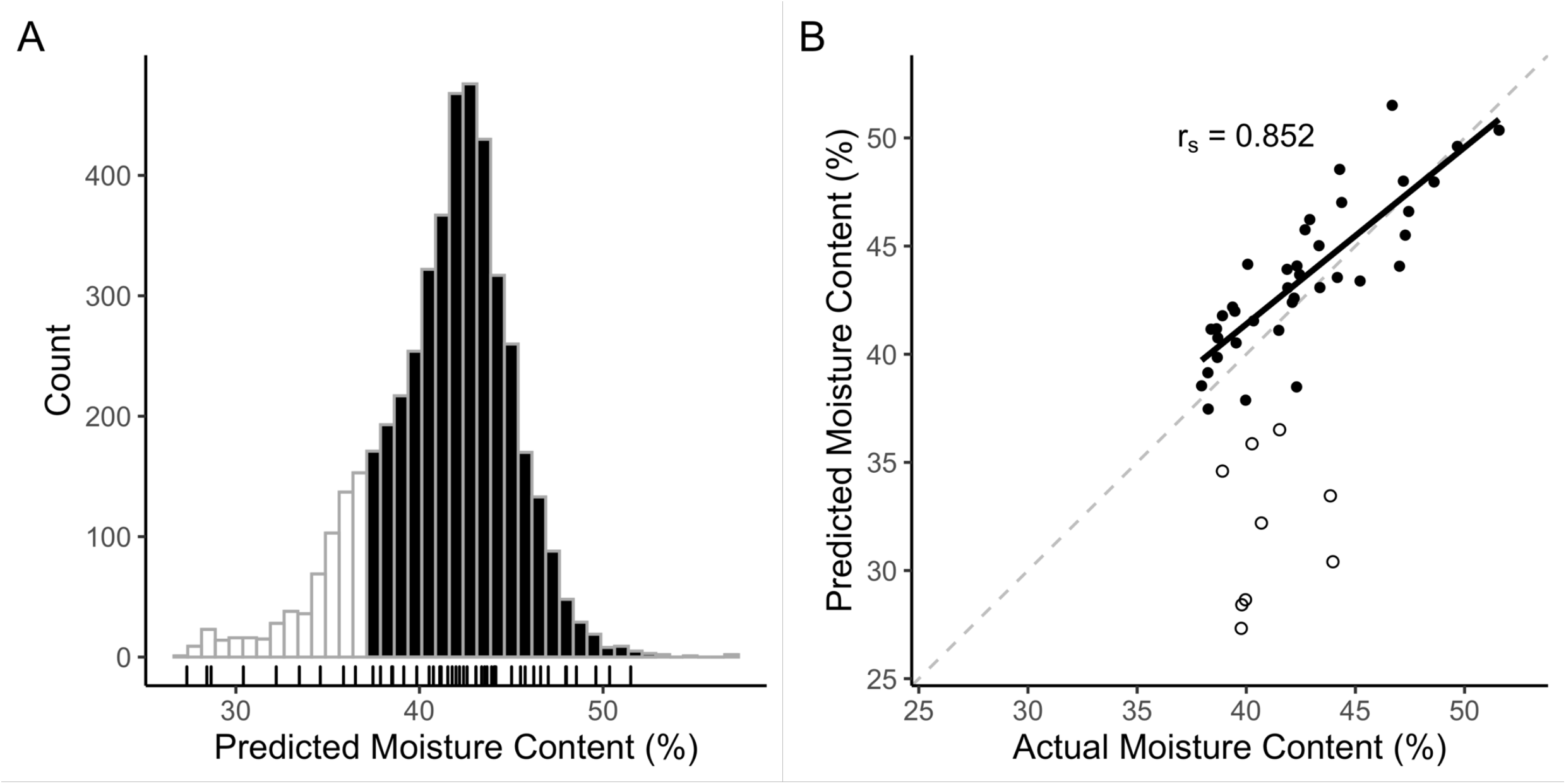
Predictions and performance of the svmLinear model. A) Distribution of predictions in the full set of 4,650 trial plots. Filled bars are within the training bounds and empty bars are extrapolated. The tick marks in the rug plot below the distribution show the 47 samples chosen for validation. B) Correlation between predicted and actual moisture content of the 47 samples chosen to validate the model. Filled circles are samples chosen within the training bounds (n=38), and empty circles were beyond the training bounds (n=9). R_s_ shown is the Spearman’s rank correlation coefficient of the filled circles. The dotted line represents a correlation of 1.0, whereas the black line shows the correlation of the filled circles.

### Understanding Factors Contributing to Features of the Final Prediction Model

To determine which wavebands have the largest influence on moisture content prediction, a permutation importance (Fisher et al. 2019) test was run on the linear SVM kernel. This was performed by collecting a baseline r^2^ value of the model trained on the full dataset predicting on the 38 validation samples within the training bounds, and comparing this to the r^2^ value calculated on the same samples after permuting each waveband in the validation samples. A larger difference between the baseline and permuted performance is indicative of a larger importance to the model. It was found that many of the spectra are roughly equally important in the prediction of moisture content (Figure 4A).

**Figure 4.**
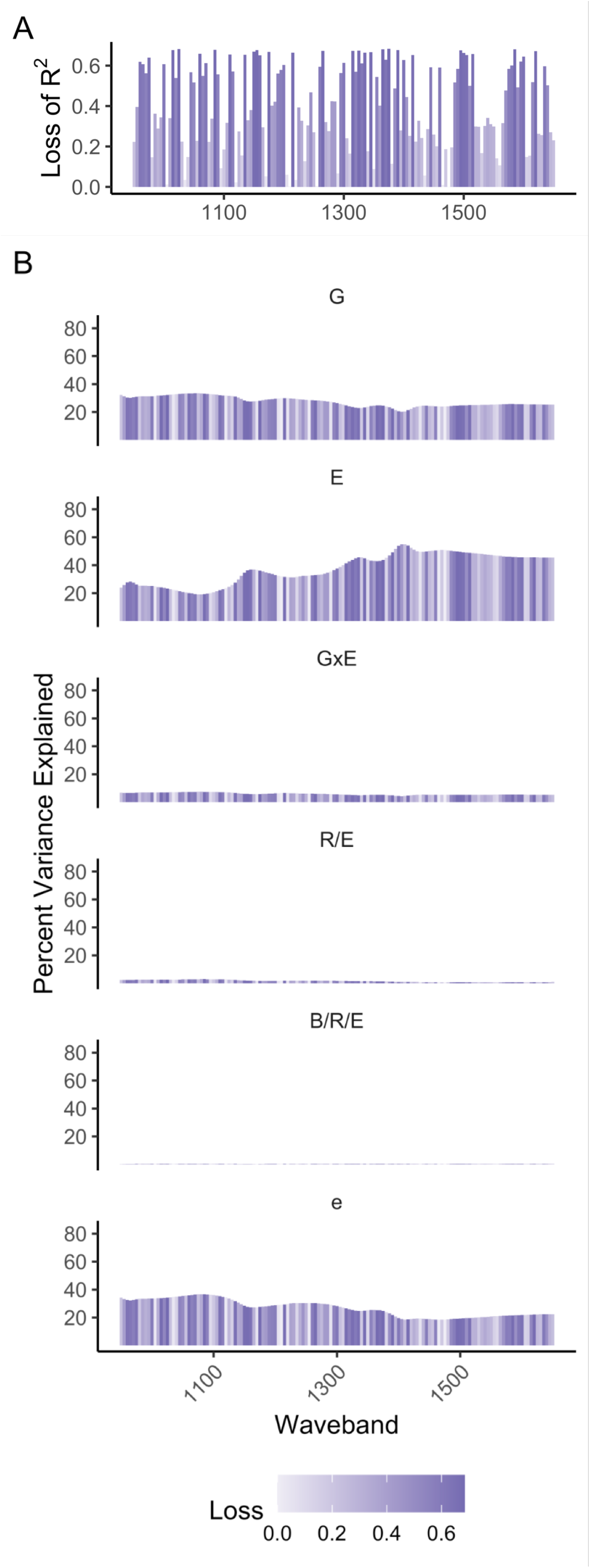
Importance of waveband for moisture content prediction and variance partitioning for wavebands. A) Importance of each waveband to the linear SVM model with height and hue equal to the loss of information when each waveband is permuted in the validation set. Values represent the average of 100 permutations for each waveband. B) Variance partitions for each waveband with height equal to variance explained, and hue equal to importance found in A. G, genotype; E, environment; GxE, genotype-by-environment interaction; R/E, replicate nested within environment; B/R/E, block nested within replicate nested within environment; e, error.

A linear mixed model was used to analyze sources of variation in the spectral data across the full set of 3,987 trial plots grown in five environments that were predicted within the boundaries of the training data. Percent variance explained by different factors in the model varied greatly across the spectral wavebands (FIgure 4B). For example, genotype explained 20.3% to 33.5% (average 27.5%) of the variation across the wavebands, and environment explained 19.1% to 55.0% (average 37.7%) of the variation. Variance explained by genotype was relatively constant across the wavebands, whereas the environment had an increasing percent variance explained as wavebands increased.

### Understanding Factors Affecting Moisture Content Predictions

To assess which sources of variation have the greatest impact on moisture content predictions from the linear SVM kernel, a random effects model (Bates et al. 2015) was created for moisture content prediction as a function of genotype, environment, genotype-by-environment interaction, and block nested within replication nested within environment, on the full set of 3,987 trial plots that were predicted within the boundaries of the training data (Figure 5A). Environment played the largest role in the prediction of moisture content, explaining 49.9% of the variation, whereas genotype and genotype-by-environment interaction only explained 16.6% and 11.8%, respectively. The three locations these trials were grown (St. Paul, MN, Boone County, IA, and Columbia, MO) are geographically diverse, and climatic conditions, especially with regards to precipitation, were very different between 2016 and 2017 for the MN, IA, and MO locations (Supplemental Figure 4; Supplemental Table 8), contributing to the large effect of environment in this experiment.

**Figure 5.**
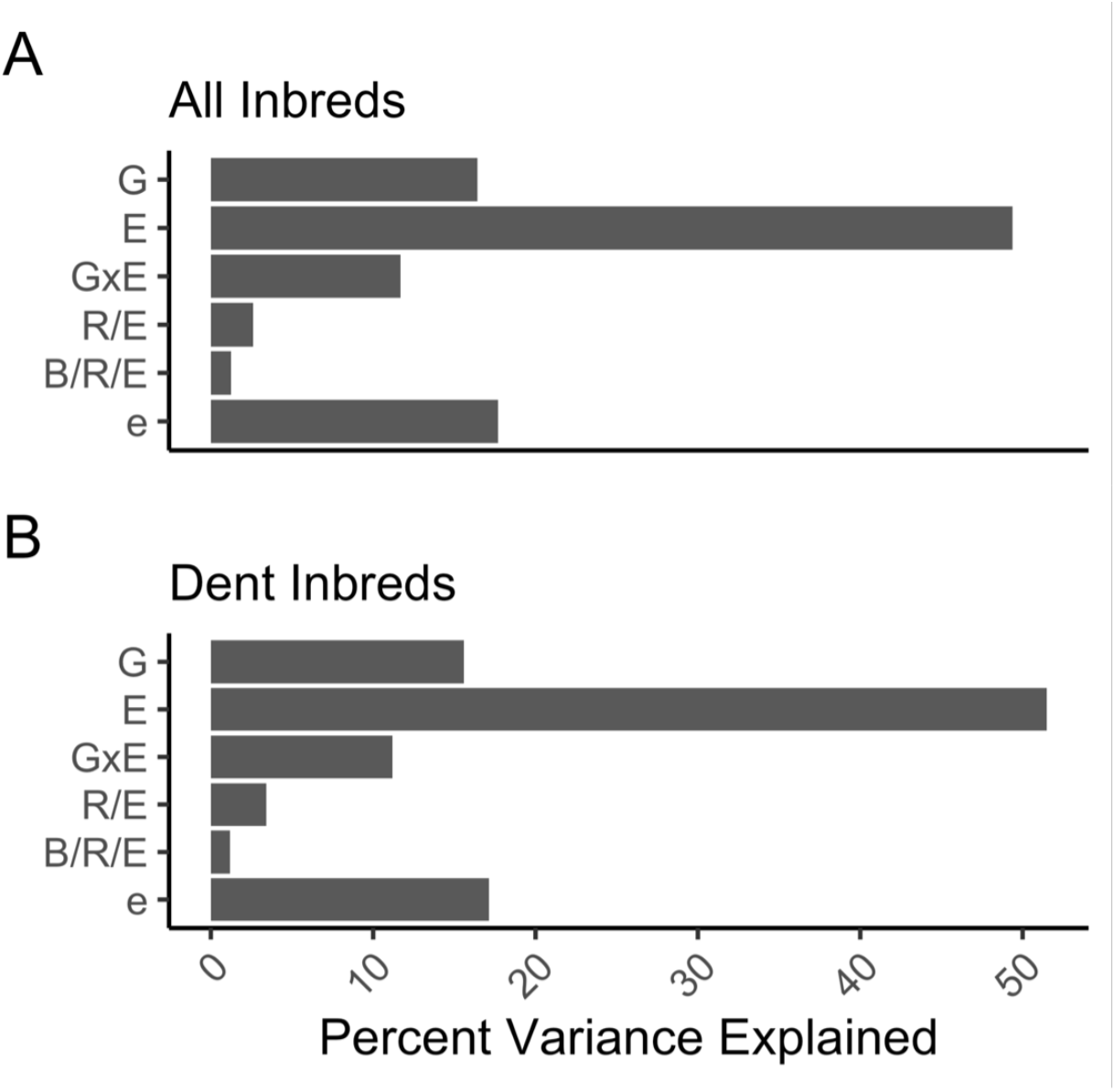
Partitioning of moisture content prediction variance. A) Variance partitioning for all inbreds in the full set of 3,987 trial plots. B) Variance partitioning for only dent types of maize (n=2,813). G, genotype; E, environment; GxE, genotype-by-environment interaction; R/E, replicate nested within environment; B/R/E, block nested within replicate nested within environment; e, error.

The majority of food grade corn grown in the United States is dent corn. As such, the same random effects model was applied to just the dent corn lines to determine if percent variance explained by the sources of variation in this subset of lines differed from the larger population of genotypes. In the subset of dent plots, variance explained by environment increased slightly to 51.5%, and variance explained by both genotype and genotype-by-environment interaction decreased to 15.6% and 11.2%, respectively (Figure 5B). Analysis of variance both in the full dataset as well as in this subset of lines support that growing environments will play a large role in the predictions of moisture content, and that environment will need to be considered when breeders test and recommend lines for improved moisture content.

The relationship between moisture content prediction and kernel macromolecular traits was tested next. Most compositional traits were significantly correlated with moisture content prediction (Supplemental Figure 5). Protein, starch, fat, and fiber were all significantly correlated (p<0.05) with predicted moisture content on the full dataset (n=3,987 plots). Protein and fat showed weak correlations of −0.099 and 0.097, respectively, while starch and fiber showed stronger correlations of −0.475 and −0.408, respectively. The exact mechanistic role of each of these compositional attributes and interactions between them to determine moisture content is still unknown, however, these data support a role for a number of compositional traits in contributing to this complex trait.

Heritability of moisture content prediction was also determined. Moisture content prediction had an entry-mean heritability of 0.80. This moderate heritability suggests that gains from selection can be made for this trait. While the heritability is moderately high, it is important to remember that the environment plays a large role in moisture content prediction, and will need to be accounted for throughout the breeding process.

### Genetic Dissection of Predicted Moisture Content

A genome wide association study (GWAS) was performed using GAPIT (Lipka et al. 2012) and FarmCPU (Liu et al. 2016) to dissect the genetic architecture controlling moisture content. Because environment explained a substantial portion of variation in this experiment, BLUPs were extracted for each environment (Supplemental Table 1). The BLUPs were normally distributed in each environment (not shown) with an average range of predicted moisture content BLUPs of 6.95 across the environments. Environments 1, 2, 4, and 5 all had ranges in variation greater than 7, while environment 3 had a very low range in variation of 1.14. The GWAS identified 26 significant SNPs across the five environments (Supplemental Table 9) at the significance threshold determined for each environment (p_MN2016_=1.04×10^-7^, p_IA2016_=6.17×10^-8^, p_MN2017_=2.23×10^-9^, p_IA2017_=2.13×10^-8^, p_MO2017_=1.41×10^-7^). Eight SNPs were found to be significantly associated with moisture content prediction in both MN 2016 and IA 2016, and five SNPs were found in both IA 2017 and MO 2017 (Figure 6), none of which were shared across the environments. The effects each SNP had on moisture content ranged from −0.71 to 0.73 (average=-0.01). While none of the significant SNPs were shared across environments, common annotations and functions across environments were observed (Supplemental Table 9). Development and stress response genes were found in each environment, though from different genes in each environment. Also noteworthy was the identification of significant SNPs within genes that encode auxin response proteins and energy metabolism proteins, each found in three of the four significant environments.

**Figure 6.**
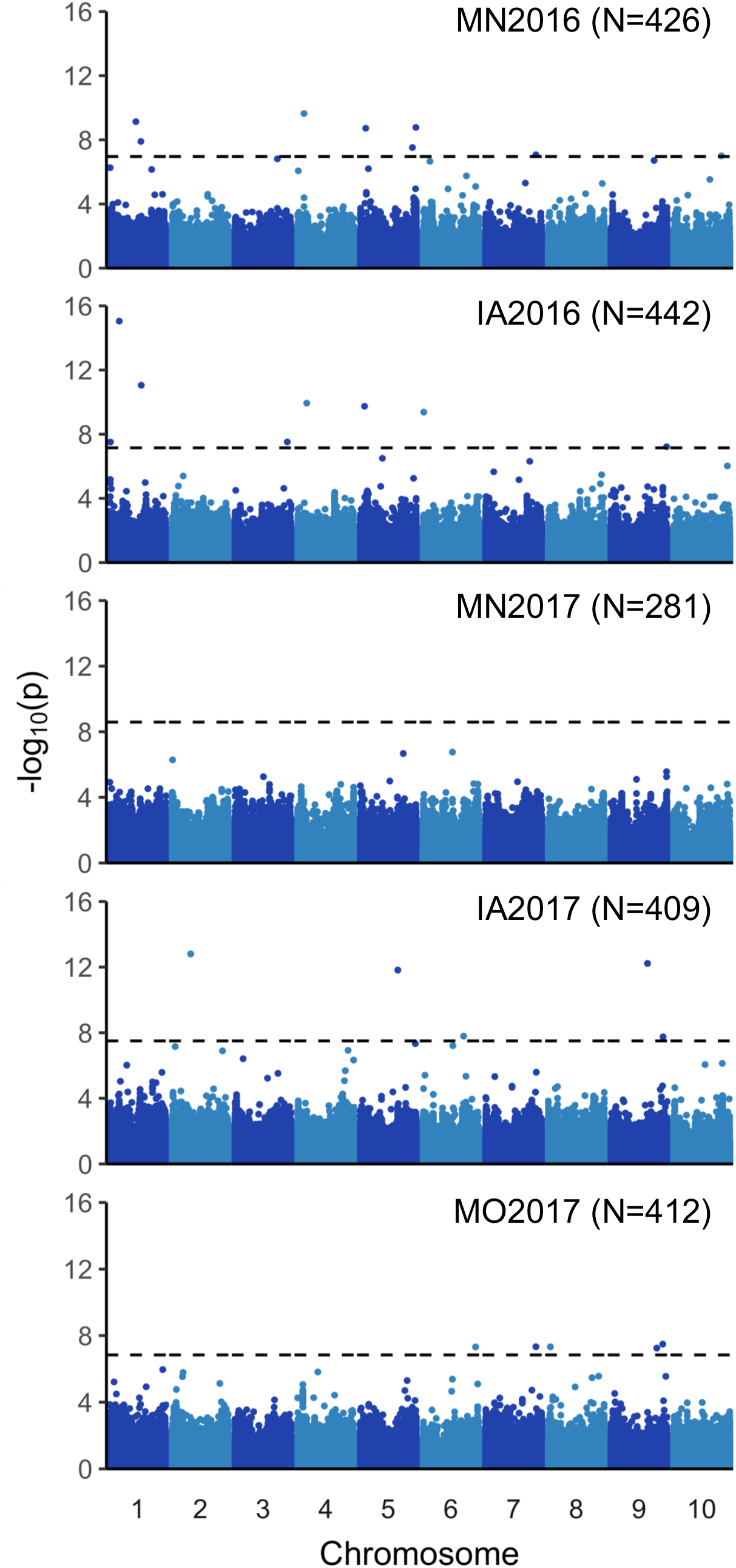
Genome-wide association study (GWAS) for moisture content prediction across environments. GWAS was performed using 2,412,791 genome-wide SNP markers on BLUPs from 281-442 genotypes in each of five environments that had sufficient yield for NIR scanning and available marker data. Dashed line shows the permuted significance threshold.

To further understand the complexity of the genetic architecture controlling moisture content, we estimated the percent variance explained by both the significant SNPs in each environment and the entire SNP dataset. Significant SNPs explained between 5.04% and 16.01% (average=10.41%) of the variation in moisture content prediction, whereas the entire SNP set explained 10.54% to 45.99% (average=31.68%) of the variation (Supplemental Table 10). This suggests that a significant amount of the genetic variation of moisture content prediction is controlled by many small effect loci.

## DISCUSSION

Current testing for kernel moisture content is done either through proxy tests (such as kernel hardness), or through pilot plant cooking trials (Holmes et al. 2019). The proxy tests are fast, but not very quantitative or precise. The pilot plant trials require a large quantity of a grain sample (500-1000 lbs) for large scale cooking, and are thus slow and low throughput, which is not amenable to the large-scale numbers and small-scale sample amounts that are common in plant breeding. The methods developed in this study are high throughput, quantitative, and optimal for early stages in the breeding process when there are many samples to test for moisture content.

In this study seven different models (lm, pls, pcr, rf, svmLinear, svmPoly, and svmRadial) were tested to see which could best predict moisture content during nixtamalization from raw kernel NIR absorbances. These models were chosen for a number of reasons. The linear model (lm) was chosen as a baseline model that all other models should outperform, the partial least squares (pls) and principal component regression (pcr) models were chosen for their dimensionality reduction capabilities (Mevik and Wehrens 2007), and the random forest (rf) and support vector machines were chosen for their general model robustness and applicability (Breiman 2001; Karatzoglou et al. 2004). The model that performed best was the linear SVM kernel. This model seems to fit the absolute value dataset very well given its high predictive capabilities with a range of hyperparameter values (Supplemental Table 7). This model performs comparably to previous work using PLS models to predict maize kernel composition that observed an average r^2^ value of 0.654 and 0.815 (Orman and Schumann 1991; Spielbauer et al. 2009). The relative performance of the linear SVM kernel over the pls model is also consistent with previous work which indicates that regression SVM kernels can outperform both PLS and artificial neural networks in complex, high-dimensional datasets such as predicting composition from NIR waveband absorbances (Balabin and Lomakina 2011).

Application of this model to a diverse set of germplasm grown in replicate in five environments revealed a significant effect of environment on observed phenotypic variation. End of season growing conditions varied substantially for the five environments used in this study (Supplemental Table 8; Supplemental Figure 4), which could impact grain filling, composition, and other grain quality factors such as kernel cracking. MO 2017 was the driest environment while also accruing the most growing degree days. Conversely, MN 2016 and IA 2016 had average rainfall events, but saw a plateau in cumulative growing degree days at the end of the season. These results highlight the importance of not only genetics, but also agronomics and growth conditions that are important in obtaining grain that meets industry standards for food grade applications.

A genome-wide association study (GWAS) was performed to determine SNPs that were significantly correlated with moisture content prediction. Twenty-six significant SNPs were found across four of the five environments used in this study. Genes related to development and stress response were found in each environment, through different genes in each environment. It makes sense that the multitude of stresses presented by each environment would differentially affect the development of the plants, changing the kernel composition (Mayer et al. 2016; Sehgal et al. 2018), and with it the NIR spectral profile of the grain samples (Spielbauer et al. 2009). The number of stress response and development genes that were significant in the GWAS, coupled with the large percent of moisture content prediction variance explained by environment, indicates that breeders should consider the stress response of a genotype when breeding for moisture content after nixtamalization. It also suggests that lines exhibiting more robust environmental stress tolerance could maintain a more consistent moisture content prediction across environments, though more testing would be needed to confirm this.

In conclusion, our linear SVM kernel trained on absolute value normalized spectra is a high throughput, quantitative and robust system for predicting moisture content in maize without cooking a single kernel. This method will assist breeders by allowing them to screen a large pool of germplasm to quickly obtain quantitative trait values for selection. Breeders can utilize this system during the early stages of breeding while the number of samples is high, and transition to pilot plant studies during later stages as sample numbers decrease. Manufacturers of masa-based products may also be interested in incorporating this system into their pipelines to predictively alter cooking conditions. Rather than sampling kernels for moisture content after nixtamalization and adjusting pH, cooking time, or steeping time post hoc, they could instead scan samples with an NIR before cooking to predict the moisture content of the kernels, and adjust the cooking conditions accordingly.

## ACKNOWLEDGEMENTS

The authors acknowledge the Minnesota Supercomputing Institute (MSI) at the University of Minnesota for providing resources that contributed to the research results reported in this paper.

## DECLARATIONS

### Availability of data and material

All raw data is included in the supplemental tables.

### Code availability

All code is publicly available on GitHub at https://github.com/HirschLabUMN/ML_Moisture_Prediction.

### Author contributions

CNH, GA, MYN, SFG, DE, AW, NA conceived this experiment. MJB, JSR, AMG, TJH, MH conducted the experiments. MJB, JSR, DPE, analyzed the data. MJB visualized the data. MJB and CNH wrote the original draft. All co-authors edited and approved the final manuscript.

### Ethics approval

Not applicable

### Consent to participate

Not applicable

### Consent for publication

Not applicable

---

**Supplemental Figure 1.**
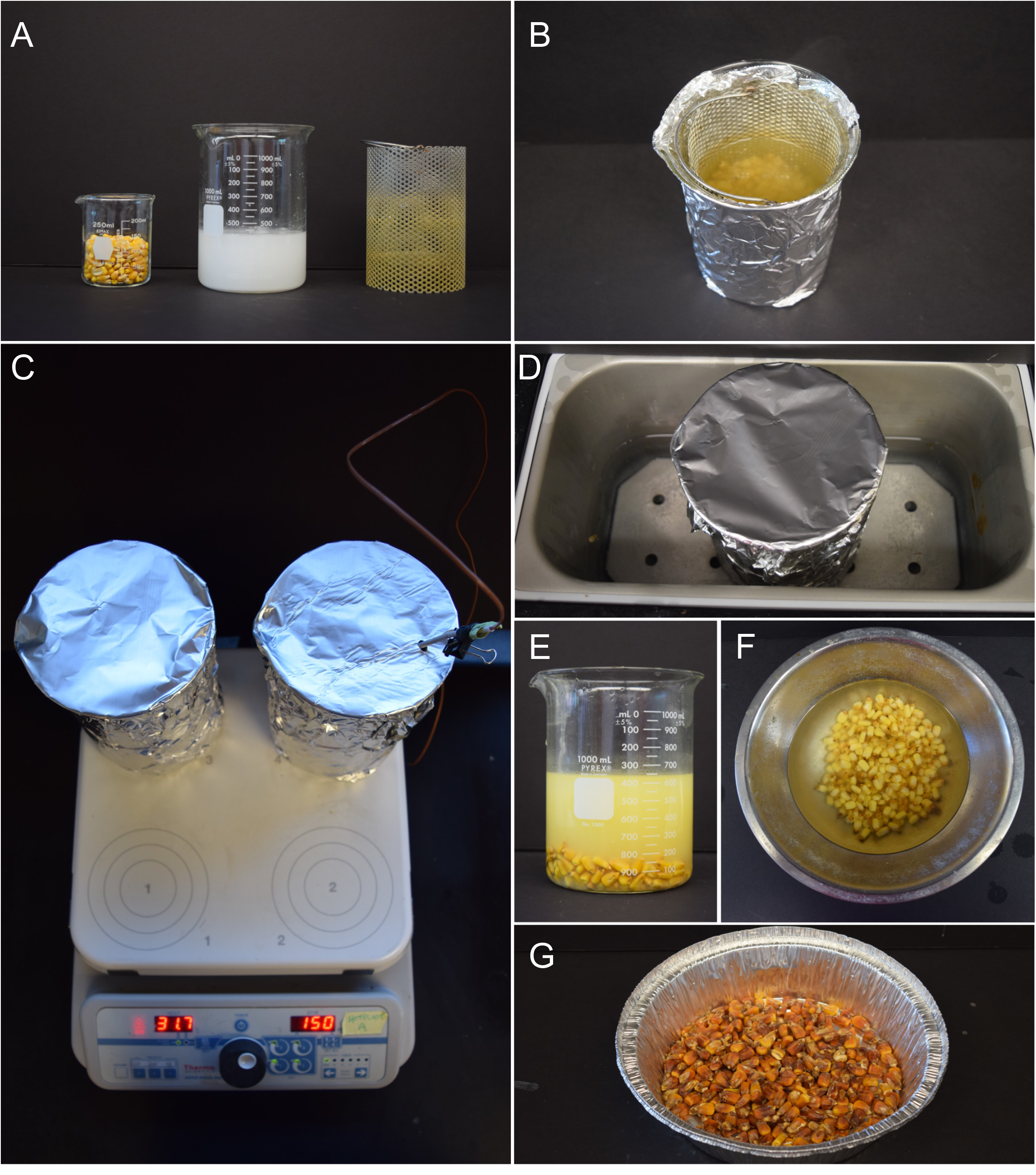
Steps in the scaled-down benchtop cook test. A) Supplies needed to cook one sample (stir bar not pictured): from left to right, 100 g dry sample, 2.4g of calcium hydroxide and 400ml of deionized water, and cooking basket. B) Tinfoil wrapped beaker ready to be placed on the hotplate. C) Two samples being cooked on the hotplate, upper right position 4 monitors the temperature. D) Sample in a 49°C water bath. E) Sample after steeping; leached solids are suspended in nejayote. F) Kernels during pericarp removal by hand. G) Kernels that were dried in a 103°C oven to determine final dry kernel weight.

**Supplemental Figure 2.**
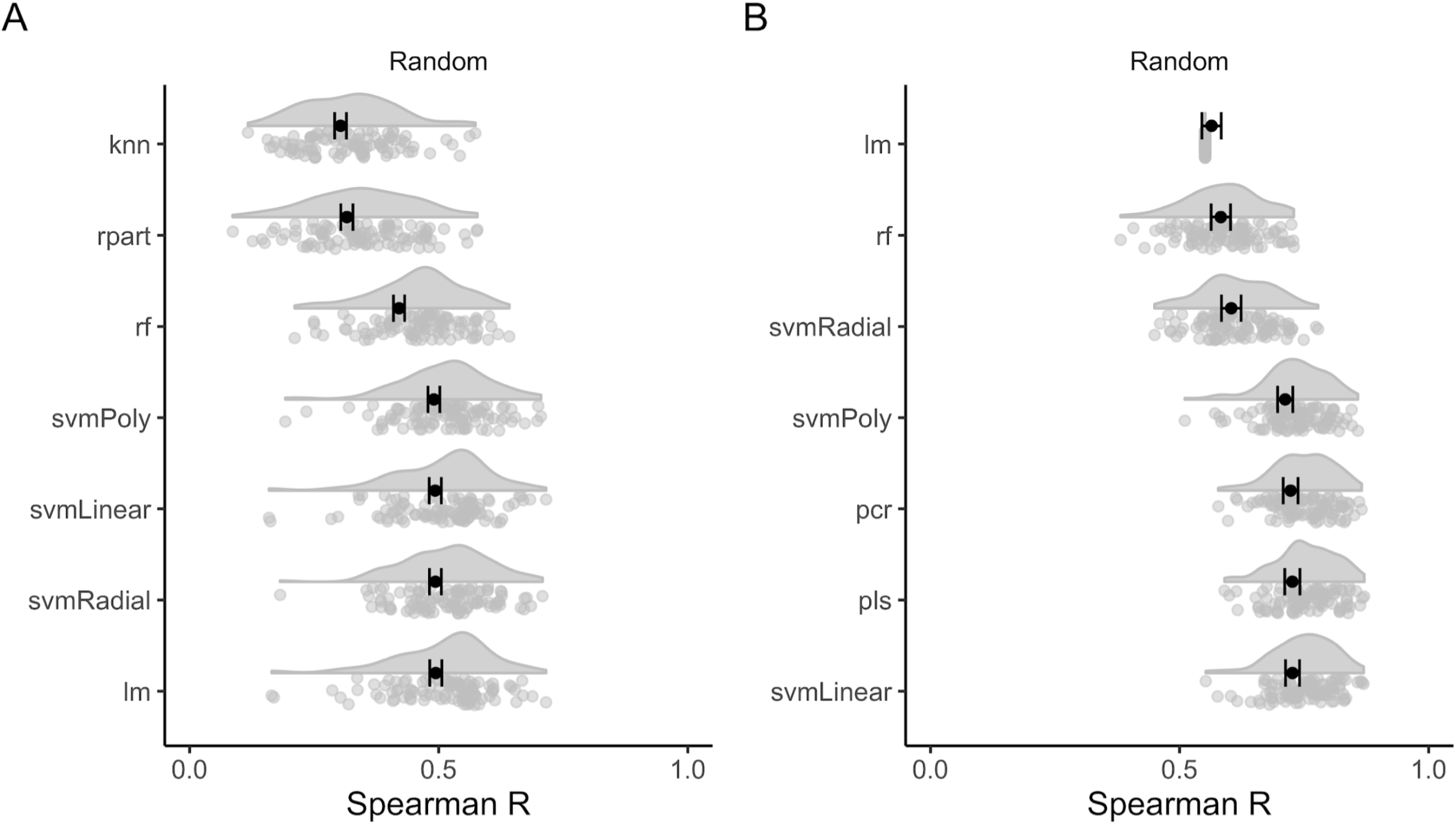
Bootstrapped performance of top macromolecular and spectra models partitioned and cross validated randomly. A) Bootstrapped performance of the top macromolecular models. B) Bootstrapped performance of the top spectra models.

**Supplemental Figure 3.**
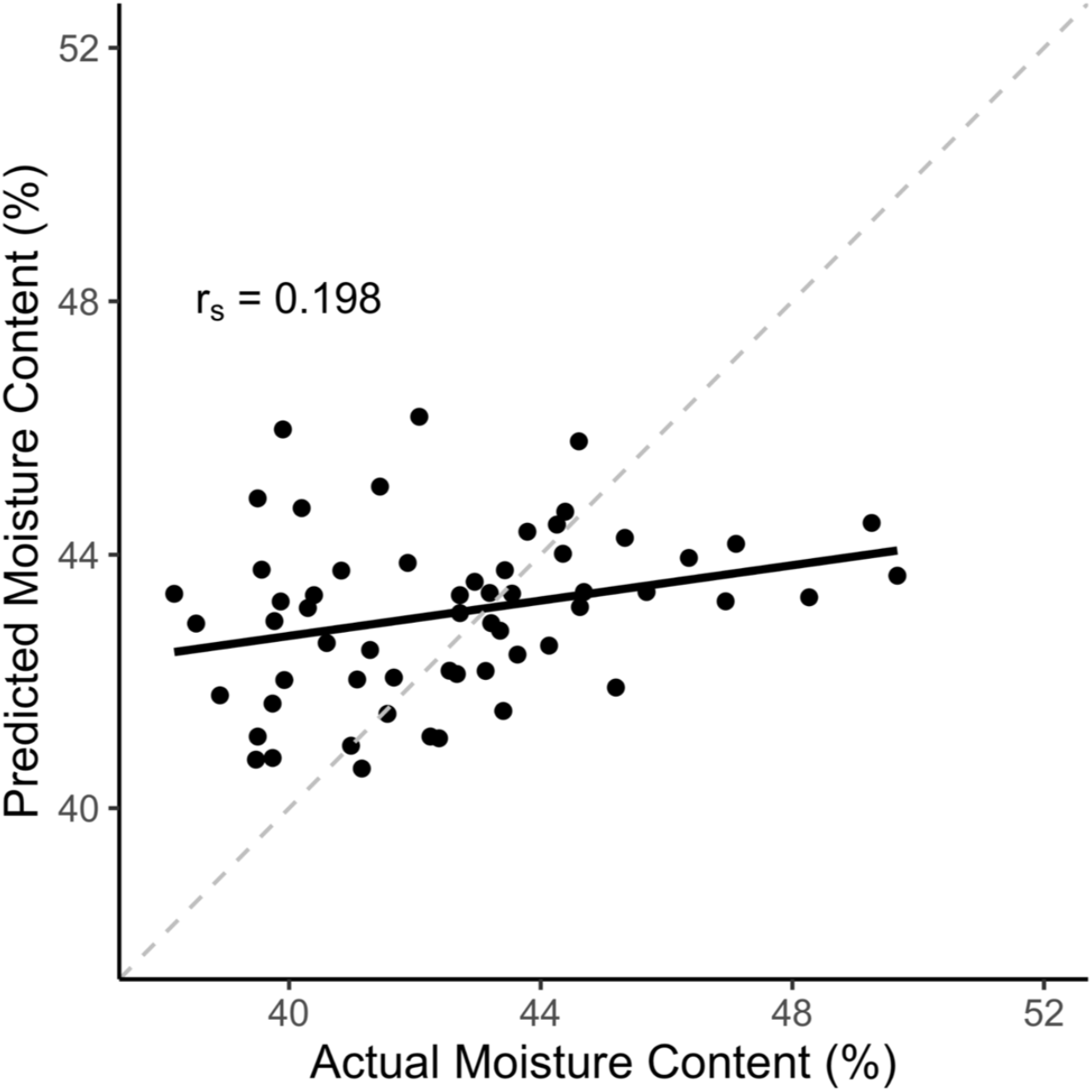
Example of a bootstrapped performance. This plot shows the performance of a macromolecular KNN model with a k value of 5, partitioned and cross validated by genotype. The data points shown are a 20% subset (n=63) of the full 316 samples, which were partitioned out to determine performance of the model. This was done 100 times for each model, hyperparameter, partitioning and cross validation type, and normalization (in the case of spectra models) combination to reduce the effect of randomness (Figure 2 and Supplemental Figure 2). The dotted line in this image shows a 1:1 ratio, and the black line shows the correlation of the model’s predictions.

**Supplemental Figure 4.**
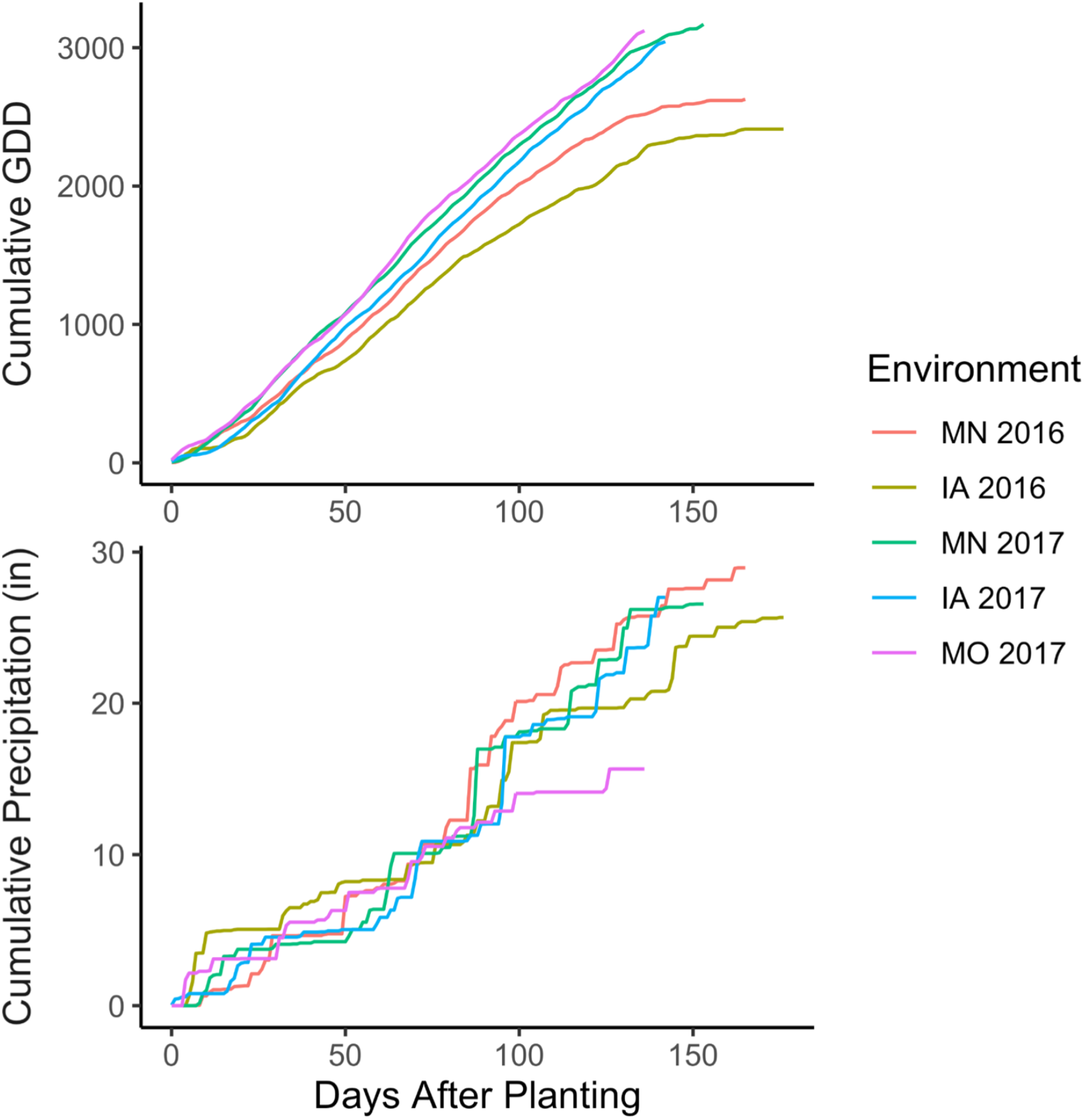
Cumulative Growing Degree Days and Precipitation Across Environments. This plot shows the environmental differences in cumulative growing degree days (GDD) and cumulative precipitation. All plantings occurred between May 8 and May 17, and all harvest occurred between September 27 and October 24.

**Supplemental Figure 5.**
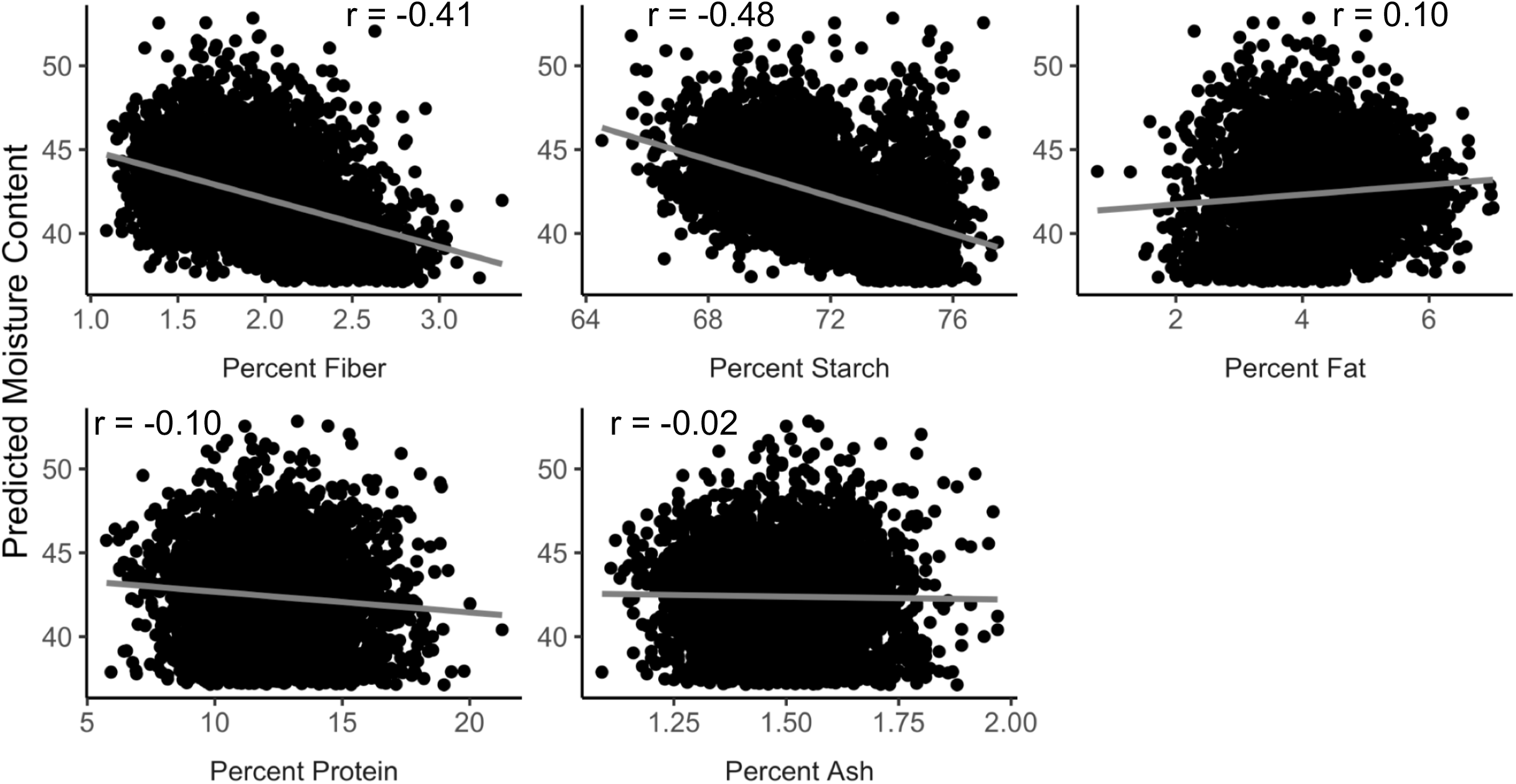
Correlation between compositional traits and moisture content. Data are for predictions from 3,987 plots grown over five environments. Macromolecular trait correlated to moisture content prediction is shown beneath each plot. R value shown is the pearson correlation coefficient. All correlations were significant at p<0.05 except ash content.

